# mt.surv: Multi-Threshold Survival Analysis for Associating Continuous Predictor Variables with Time-to-Event Outcomes

**DOI:** 10.1101/2025.05.29.656703

**Authors:** Alexander Loncar, Rebecca Hoyd, Yunzhou Liu, Caroline Dravillas, Dipankor Dhrubo, Daniel Spakowicz

## Abstract

**Motivation:** Time-to-event models are a useful and common approach to infer the importance of biological variables. Most often, the predictor variable is binarized and associated with censored time to death—a so-called Kaplan-Meier curve. However, the threshold to binarize the continuous variable is often arbitrary. We sought a rigorous way to define thresholds and evaluate the strength and consistency of association with the event of interest.

**Results:** We present {mt.surv}, an R package for multi-threshold survival analyses. The primary function performs a time-to-event analysis, where a continuous predictor variable is stratified into two groups at an arbitrary number of places, defined by the percentile of the distribution. The result can be visualized with a function that creates a custom line plot with the -log of a log-likelihood p-value against the threshold percentile. A third function operates on this plot and calculates the area above a significance threshold, creating a scalar that can be used to rank biological variables for their association with the event of interest. This framework can be broadly applied to any continuous predictor variable including, but not limited to, gene expression and microbial abundances. Several helper functions are included for structuring input data and job submission in a cluster framework. We found that this method has value in discovery-type analyses where one lacks prior information about appropriate stratification thresholds. However, we found additional biological insight is possible— indeed, quite common—as many variables show different associations at different stratification thresholds.

**Availability:** On CRAN as {mt.surv}.

## 1 Introduction

Time-to-event models are widely employed to evaluate the relationship between biological variables and clinical outcomes over time. For example, in randomized controlled trials (RCTs), the hazard ratio estimator serves as a foundational tool for assessing survival probability, accounting for both event occurrence and censoring to ensure unbiased estimates even with incomplete follow-up (Greenhouse et al., 1989). Despite their importance, there are a wide variety of approaches used to estimate risk from continuous predictor variables.

Common practice in time-to-event models includes simplifying continuous predictor variables into a binary format, such as high versus low expression levels of a gene or treatment versus control groups, to assess their association with censored outcomes. This approach facilitates the visualization of outcomes over time, such as survival probabilities, using Kaplan–Meier (K-M) curves, a non-parametric method that estimates the survival function from observed survival times, accounting for right-censored data (Kaplan & and Meier, 1958). The log-rank test is often used in conjunction to determine statistical significance between the stratified survival curves, providing insight into the prognostic value of the binary variable (Bland & Altman, 2004). While this dichotomization simplifies interpretation, it may lead to a loss of information and statistical power compared to continuous modeling (Royston et al., 2006). On the other hand, continuous variables may exhibit nonlinear relationships with outcomes, varying across different segments of the predictor’s distribution, which challenges the assumption of linearity often made in basic regression models. This behavior implies that the risk does not increase or decrease at a constant rate across the distribution of the variable. Modeling strategies, such as restricted cubic splines (Pichardo Rodriguez et al., 2024), fractional polynomials, and non-linear mixed effects models (Nguyen Phuong et al., 2024; Royston et al., 2006), have been explored to more accurately capture these relationships, thereby avoiding the information loss and bias that result from simple binarization or linear assumptions.

Here, we introduce {mt.surv}, an R package that supports exploratory analyses of the relationship between a continuous variable and a censored time-to-event outcome. Included are functions for estimating the association with outcome at multiple binary thresholds and then quantifying the strength of the relationship with a novel scalar “area”, a cardinal value that can be compared between datasets. The package is broadly applicable to any continuous data (gene expression, microbial abundance, etc.) and has particular utility when the variable shows different behavior at different parts of its distribution.

## 2 Methods

### 2.1.1 Overview and Implementation of the Package

We developed the mt.surv R package to perform survival analysis systematically splitting the dataset into various thresholds in a continuous pattern. This enables users to evaluate the statistical correlation with survival outcomes by dividing their dataset into two groups at each level. Identifying the optimal cut points in any count data that categorizes patients into high-risk and low-risk groups is one of the package’s primary applications. Key functions include (i) survivalByQuantile(), which generates a table with cox analysis parameters at various threshold quantiles, and (ii) calculateArea(), a function that integrates the area above a 0.05 log likelihood p-value threshold and outputs a scalar that can be used to rank the importance of different features (**Figure 1**). This package provides the optimal stratification cut point for binarizing the data and comparing it for downstream analyses.

**Figure 1.**
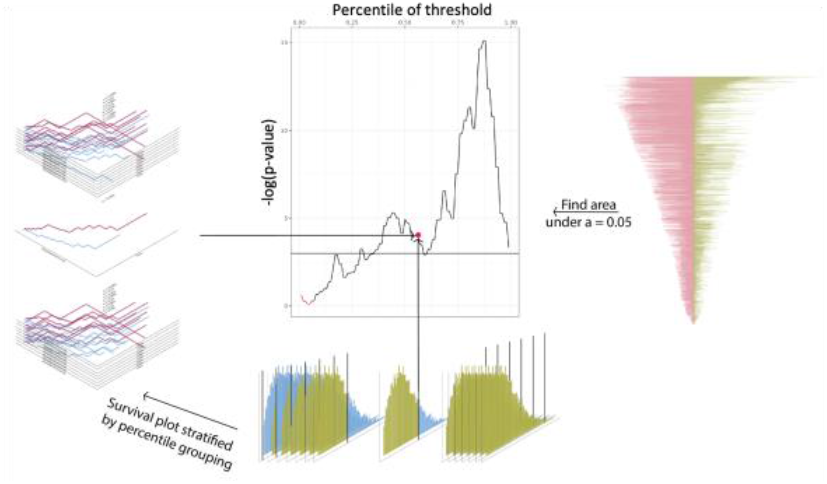
Multi-threshold survival analysis for establishing appropriate binarization of count data. A) Schematic for the process of the {mt.surv} multi-threshold survival analysis. Samples are stratified by the “high” vs. “low” abundance of each data point at every quantile from 0.01 to 0.99 (i.e., quantile 0.5 represents a stratification at the median relative abundance of the count value). A survival curve is generated for each stratification, and the p-value of the survival difference in a Cox Proportional Hazards model is plotted against the quantile of the stratification. Further analysis includes the AUC beyond the p-value threshold at alpha 0.05, which is used to rank significance.

### 2.2 Package Functions

The package features several different functions conducive to the successful analysis of data. The primary functions contained in the package are as follows:

#### 2.2.1 Description of Functions Contained

##### (1) survivalByQuantile()

a. The survivalByQuantile() function computes a cox model p-value under various thresholds. The threshold is used to split the data into two distinct groups for statistical analysis. The function generates a data frame containing the hazard ratio, lower bound, upper bound, percentile, cutoff threshold, and p-value.

##### (2) calculateArea()

a. This function calculates the distance between the cutoff point p-value and p-value 0.05 as an indicator for the area under the curve. This information can be used to infer the magnitude of significance for the factor of interest. This calculateArea() function requires output from survivalByQuantile() function.

#### 2.2.2 Usage and Reproducibility

The {mt.surv} package is currently available as a development version on GitHub and CRAN.

## 3 Results

### 3.1 Application of the package

This package was applied to bulk tumor RNAseq data from The Cancer Genome Atlas to recapitulate a finding that high *Candida* tumor burden is associated with worse survival in GI cancer types (Dohlman et al., 2022). While the result validated the previous finding, testing thresholds at different percentiles of the distribution using {mt.surv} revealed the opposite behavior at low *Candida* abundances. This exemplifies the need for a standardized method that {mt.surv} provides.

In conclusion, we developed {mt.surv}, an R package that serves to infer the importance of biological variables. By stratifying continuous variables at every whole quantile, appropriate consideration of the dataset can be made.

